# Population dynamic population delineation in North American birds

**DOI:** 10.1101/2023.08.29.555290

**Authors:** Lars Witting

## Abstract

Andrewartha and Birch (1954) envisioned that natural populations are surrounded by ecological barriers, yet statistical estimates of population structure are almost completely driven by genetic analyses void of ecological information. I develop an ecological clustering routine that delineates natural populations from the synchronisation and desynchronisation in the population dynamics of continuously distributed species. The method is applied to the North American Breeding Bird Survey (Sauer et al. 2017), where it identifies two to five populations per species for 160 species, and leaves the population differentiation unresolved for 139 species. This provides one of the first statistical methods for population dynamics delineated population structure.

## 1 Introduction

The unit of population dynamics is a group of individuals of the same species that live in an area interbreeding over time. While this population concert seems simple in principle, it is vaguely defined allowing for several interpretations (Andrewartha and Birch 1954, 1984; Camus and Lima 2002; Berryman 2002). At the evolutionary level, populations are so isolated from one another that they can evolve in different directions having their own population genetic composition. At population dynamic and demographic levels, there can be a larger inter-change of individuals allowing similar genetics for different populations; yet the inter-change is so limited that each population is dominated by its own demographic and population dynamic processes. And at the local level, a population can be an aggregation of breeders in a local area, with year-to-year variation reflecting not only demography but also emigration and immigration.

While these population level concepts have been around for 3/4s of a century, the development of empirical methods for population delineation has so far progressed almost exclusively in the evolutionary field of population genetics. This makes the identification of population dynamic and demographic units a rather arbitrary process in many cases. The management of fisheries and hunts, e.g., is based on population dynamic principles and modelling, yet the identification of stocks to preserve is often based on genetics, if not simply reflecting the geographical distributions of the different fisheries and hunts. In other cases, the units to conserve might be considered national entities, or state entities within nations, with no relation to the underlying biology.

Until recently there has been an almost complete absence of statistical methods to identify demographic populations across large geographical areas (Camus and Lima 2002; Jones et al. 2007). An exception is Rushing et al. (2016) that used mean abundance and trend estimates from the North American Breeding Bird Survey (Sauer et al. 2017) to delineate populations for eight species of passerines. Applying cluster analyses to a multivariable distance matrix incorporating mean abundance, trend, and geographical position, they estimated eight to twenty populations per species.

Where trend is a crude measure of the dynamics of a population, mean abundance may reflect average demographic production, yet it is not a population dynamic measure. With the present study I develop a statistical clustering routine with the specific aim of delineating the population dynamics of continuously distributed species into geographically separated populations with separate population dynamic trajectories. The method is applied to the North American Breeding Bird Survey, where birds are counted locally at counting routes that are distributed across the North American continent, and it is possible to estimate timeseries of relative abundance for chosen areas of similar or different sizes.

The clustering method operates on a geographical grid, with separate population dynamic timeseries for the subset of sub-areas with positive counts of the species in question. While all the timeseries of point estimates of the different sub-areas are somewhat unique, this variation reflects local abundance variation and observation uncertainty at the counting routes that are integrated into the population dynamic timeseries. The timeseries are best, but uncertain, estimates of the actual population trajectories in the chosen areas.

To delineate these timeseries into trajectories of more or less independent populations, I develop a likelihood routine that clusters the strongest correlated dynamics of neighbouring areas, starting from the available local optima in the strengths of dynamic correlation across the continent. The method uses the simple principle that the spatial geographical synchronisation—and thus correlation—of population dynamics is strongest in the areas that drive the dynamics, with synchronisation and correlation deteriorating at the boundaries where the somewhat uncorrelated dynamics of different populations meet. The method is a relative method that operates on the available geographical synchronisation and desynchronisation of the dynamics, aiming to estimate the trajectories of population dynamic clusters of North American birds.

## 2 Data

The North American Breeding Bird Survey (BBS) have used skilled birders to count breeding birds by a standardised method across the states of USA and Canada every year since 1966 (Sauer et al. 2017). The survey is conducted by volunteers, and it is one of—if not the—larges systematic monitoring of wildlife on Earth, when measures by the joint scale of the area covered, the number of species involved, and the number of years with continued counting. The 2020 dataset that I analyse contain observations of 664 species/subspecies on a total of 5,210 survey routes across 62 states in North America, with approximately 2,500 routes being surveyed each year (Fig. 1).

**Figure 1:**
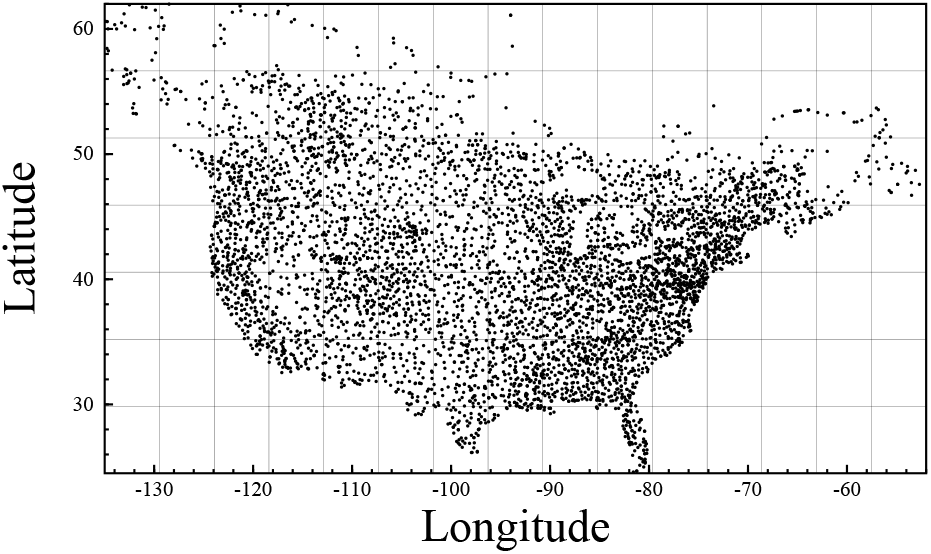
Spatial model. The geographical distribution of the study, with the survey routes of the North American Breeding Bird Survey shown by dots, and the grid of the spatial model shown by lines.

The BBS have been analysed in many ways for a variety of purposes (e.g., Robbins et al. 1986; Link and Sauer 1998; Link and Sauer 2002; Sauer et al. 2003; Smith et al. 2014; Niemuth et al. 2017; Hudson et al. 2017; Rosenberg et al. 2017; Sauer et al. 2017; Link et al. 2020). I combine traditional index estimation with a maximum likelihood based spatial clustering routine to delineate the dynamics of each species into populations with different trajectories.

To cover the majority of the BBS survey routes I use a spatial model with a 15*x*7 longitudinal/latitudinal grid (Fig. 1; excluding only the North with few routes). I use the BBS that cover this area from 1966 up to and including 2019. The data are organised in different routes, with each route having 50 point-counts placed 0.8 km apart. For each point, the observer conducts a 3 min count of all the birds heard or seen within 400 m. Some of the counts are marked as unreliable due to, e.g., bad weather, and I use only the reliable counts.

### 2.1 Index time-series

To estimate index time-series of relative density (relative number of individuals per unit area) for the different populations, I divided routes between the different observers to eliminate effects from variation in observation skills (Sauer et al. 1994). For each observer specific route, I removed the first year of the time-series because the observation efficiency is often reduced in that year (Kendall et al. 1996). I include zero observations; however, I exclude an observer specific time-series of a specific species if the observer had observed that species in less than 30% of the years in the time-series. Based on a total of 6.18 mill. observations of at least one individual of one of the 664 species/subspecies observed during the BBS, this generated a set of 18,700 observer specific time-series with observations of a multitude of species. These time-series were then used to calculate the index time-series of relative population densities for different areas.

For a species in a given area, I used the *n*_*r,t*_ = 1+ Σ *n*_*r,p,t*_ sum of the 50 point counts (subscript *p*) per year (subscript *t*) as a relative population density measure for an observer specific route (*r*), with the time-series of relative density covering all years with point counts by the observer on the route. These route counts were combined to produce index series of relative densities for each of the grid-squares where a species was observed, as well as for larger areas containing multiple neighbouring grid-squares, including the whole North America as the largest population.

The index time-series were calculated to reflect the development in the geometric mean of the relative poulation density across all the routes in an area, with the developments of relative density being calculated separately for each route. For this let

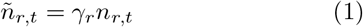

be the relative density of route *r* in year *t*, with

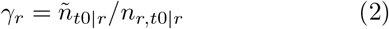

being the multiplicative scaling parameter for route *r* with *t*0|*r* being the first year with a positive count of the species on route *r* (zero counts prior to *t*0|*r* did not affect the overall index), and *ñ*_*t*0|*r*_ being the overall index estimate in that year, with the index value for the initial year being *ñ*_*t*0_ = 1, and subsequent index estimates calculated as

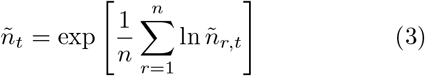

with *n* being the number of routes covered in the area in year *t*, and the error coefficient of variation of the index estimate in year *t* being calculated from the variation in the index across all the routes with data on the given species in that year. After an index series was constructed across all years, it was rescaled to make the index value of unity correspond with the geometric mean of the time-series.

The first year of an index time-series was the first year where a species was observed on at least 10 routes, and a final index series for an estimated population would not be used unless the number of observations of at least one individual was at least 30 on average across all years.

## 3 Population dynamic delineation

Having an initial set of index time-series for all the grid-squares where a species was observed, I calculated the synchronisation of the population dynamics between two population components—like grid-squares *i* and *j*—using a log-normal likelihood

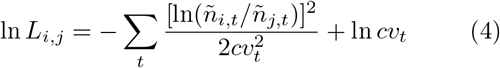

with ln *L* increasing monotonically with the synchronisation of the trajectories, *ñ*_*i,t*_ and *ñ*_*j,t*_ being the index estimates of the *i*th and *j*th population components in year *t*, and 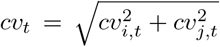 being the error coefficient of variation calculated from the coefficients of variation of both components.

I could then, for each population component *i* calculate the average population dynamic synchronisation with neighbouring population components *j*

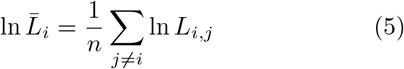

with *n* being the number of neighbouring components for population component *i*. When plotted as a map (Fig. 2, right plots), the average synchronisation identifies the local maxima where the synchronisation of the population dynamics with the neighbouring components are larger than in the surrounding areas. As evident, by comparing the left and right plots in Fig. 2, these local maxima would generally not coincide with areas of highest or lowest population density.

**Figure 2:**
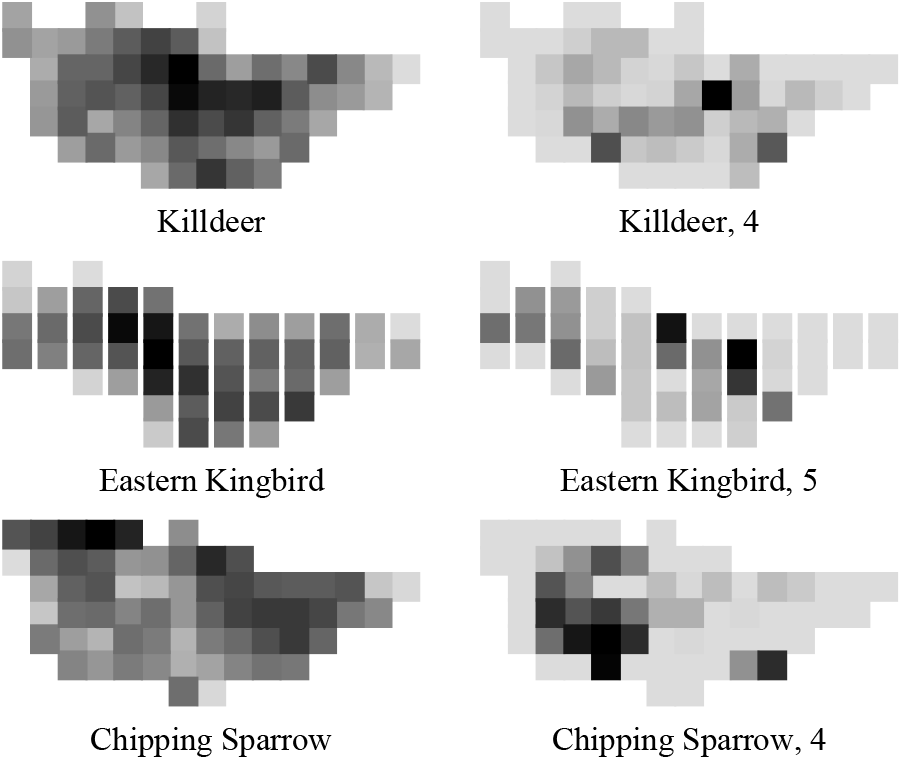
Density & synchronisation. Three cases showing the spatial distribution of relative population density (left plots; rescaled number of individuals per observation) and the local synchronisation of population dynamics (right plots; rescaled 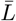), with density and synchronisation increasing with darkness. Eastern kingbird has five, and chipping sparrow and purple martin four local synchronisation maxima.

The starting points of the population dynamic clustering routine for a species, was the initial set of populations defined by the single-square local maxima populations only. I would then use the clustering routine to extend the geographical distributions of these populations until they met their natural population dynamic boundaries. For this I would for all the identified populations of a species calculate their dynamic synchronisation (eqn 4) with all of their neighbouring areas, as long as these were not already contained in an identified population. I would then identify the area with the highest synchronisation to one of the identified populations, and join the area to its neighbouring population if—and only if—the dynamics of the area was more synchronised with that population than to any of the other identified populations. If this was not the case, I would proceed with the area with the next highest level of synchronisation with a neighbouring population, until it was either possible to join a small area with a neighbouring population or there were no extra neighbouring areas available.

If it was possible to join a neighbouring area with a population, I would recalculate the index time-series for that population, and recalculate all the synchronisations between populations and their neighbouring grid-squares, and rerun the clustering routine. If on the other hand, it was impossible to find a match, I would increase the size of the non-population areas to two neighbouring grid-squares, calculate the index series of these two-grid-square components, and rerun the clustering procedure with the two-grid-square components. This increase in the number of grid-squares in the neighbouring components would continue until it was possible to join a multi-grid-square component with a population, or until it was no longer possible to find a neighbouring component with the desired number of grid-squares. If the latter was the case, the clustering routine would stop. If on the other hand, it was possible to join a multi-grid-square component with one of the identified populations, I would do that and reset the clustering routine to a comparison between single grid-square neighbours and the identified populations, continuing the process until it was no longer possible to find a single or multi-grid-square component to include into one of the identified populations.

This clustering routine produces at set of single/multi-grid-square populations from the local synchronisation maxima on the geographical grid of areas, with the boundaries between the populations being the areas with lowest population dynamic synchronisation. Yet, as the identification of the populations and their boundaries is driven by relative differences in synchronisation, the population dynamics of the identified populations could be quite similar; with the identified structuring of some species being a relatively large set of populations with quite similar. As a conservative measure, I joined neighbouring population units where the correlation coefficient between the dynamics of the two units were larger than 0.7.

A potential problem for the clustering routine relates to populations that use two or more neighbouring grid-squares dependent upon yearly fluctuations in the environment. This will induce some negative correlation between the index estimates of the grid-squares, and this might—at least potentially—prevent the relevant grid-squares from being identified as a single population dynamic cluster. To minimise this effect of year-to-year fluctuations, and to focus more on the underlying factors that control the overall shapes of the population dynamic trajectories, I used the five-year running mean of the index time-series in the statistical calculations of synchronisation and correlation. The final index series were the raw series with no smoothing.

### 3.1 Spatial resolution

An issue is the spatial resolution of the data. If the underlying grid has too small grid-squares, the estimated index series of the different squares would be so noisy that we cannot trust the estimated delineation. The clustering routine would then tend to identify too many populations where the differences in the estimated trajectories reflect uncertainty rather than differences in the underlying dynamics. If on the other hand the grid-squares are too large, the spatial resolution would be too small to differentiate populations.

To identify an optimal grid size for the clustering routine, I used the average squared residual

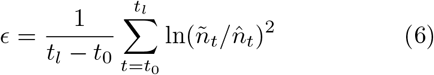

between an index series (*ñ*_*t*_) and its five-year running mean 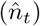 as an estimate of the uncertainty in the estimated trajectories, considering both observation and process error. I could then for different grid sizes calculate the average 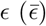 across the index series that were estimated for all the populations that were identified by the clustering routing across all species. I then minimised 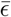 by changing the grid size to find that the most accurate (i.e., least year-to-year fluctuating) index series were estimated by a 15*x*7 longitudinal/latitudinal grid (Fig. 3).

**Figure 3:**
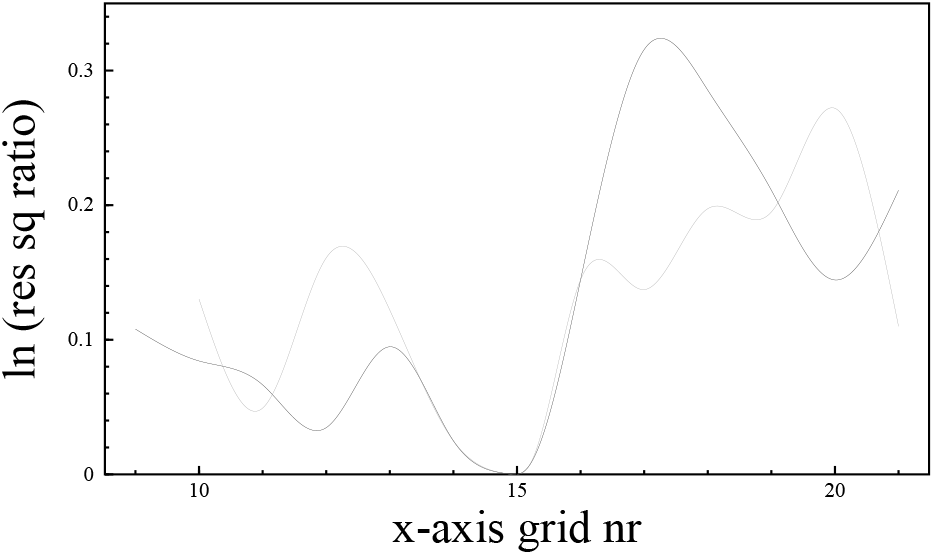
Spatial resolution. Cubic spline interpolation curves for the natural logarithm of the 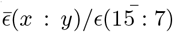-ratio for the average squared residual across all estimated populations, with *x* : *y* being the grid size of the spatial model, with the most accurate index series following form the 15 : 7-grid in Fig. 1. The black curve reflects a *x* : *y* = *x* : *x/*2-grid, and the grey curve a *x* : *y* = *x* : 7-grid.

## 4 Results

A total of 626 local maxima (ln 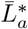) in the synchronisation of population dynamics were identified across 299 species of North American birds that passed the minimum data criterion for population delineation.

The population dynamic clustering routine failed to identify explicit geographically distributed populations for 139 species; usually because there was only a single local synchronisation maximum. For cases with two or more local maxima, the routine identified a single population for 60 species, two populations for 59 species, three populations for 24 species, four populations for 12 species, and five populations for five species.

In all I obtained 323 population level time-series with identified boundaries, and 139 time-series at the level of North America with no explicitly identified boundaries. The geographical distributions and index time-series for forty species are plotted in Fig. 4, with the population delineations of all species plotted in Supplementary Information Si1. All estimated time-series are listed in Supplementary Information Si2.

**Figure 4:**
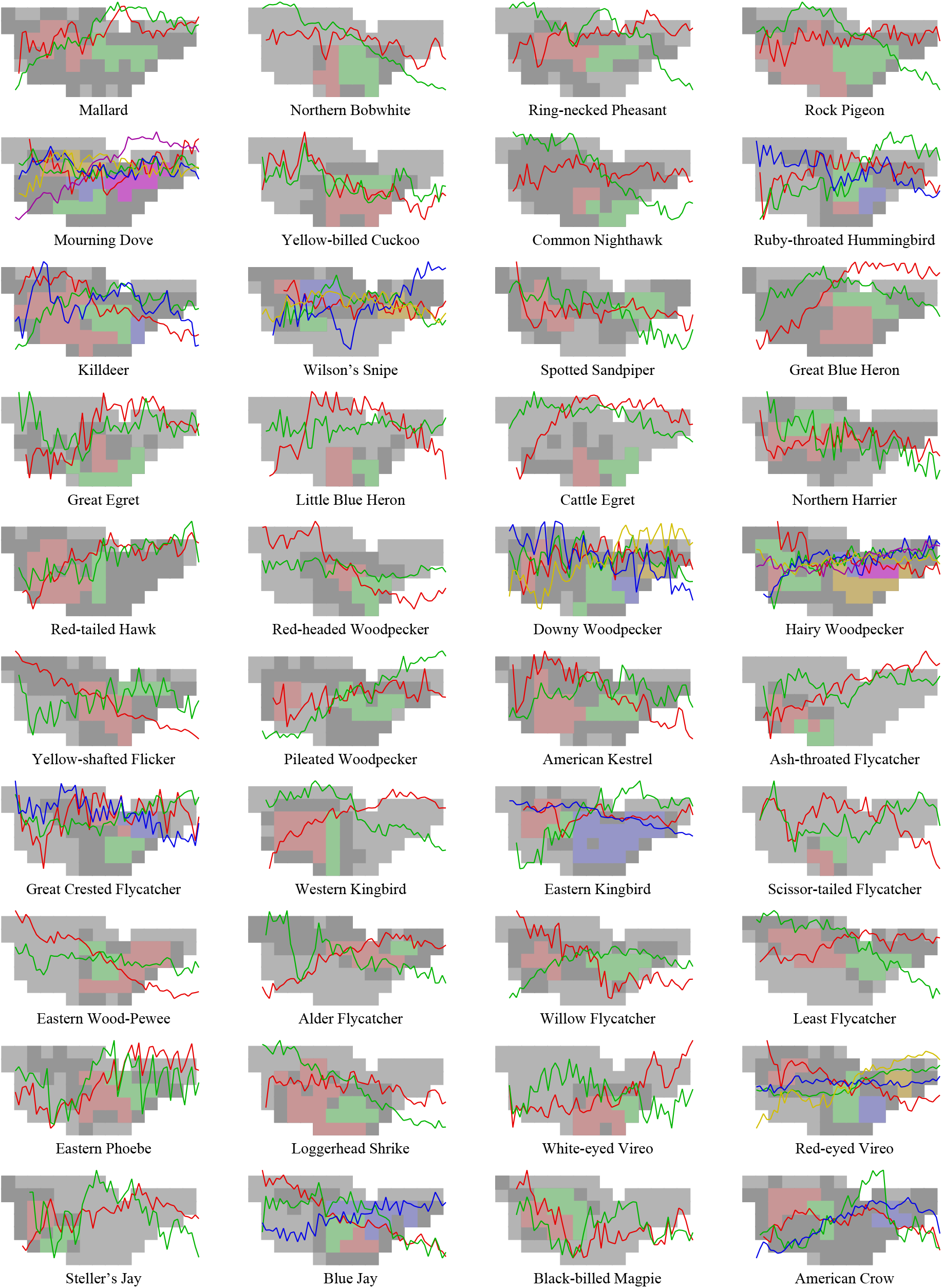
Population delineation in forty species as estimated by the clustering routine, with colours separating the distributions and index trajectories of the different populations. Dark grey mark areas that were not selected by the clustering routine or selected but with less than 50 observations per year. All delineations are plotted in Supplement S1, with the estimated index trajectories listed Supplement S2 and S3.

## 5 Discussion

The Moran effect (Moran 1953; Ranta et al. 1995) refers to cases where spatially correlated fluctuations in density-independent factors synchronise population dynamic fluctuations over larger areas. Other factors that can synchronise the spatial dynamics of a species includes trophic interactions (Ims and Andreassen 2000) as well as dispersal across locally regulated populations (Bjørnstad et al. 1999; Kendall et al. 2000). Spatially synchronised dynamics have been widely confirmed for a variety of taxa, including insects (Hanski and Woiwod 1993; Sutcliffe et al. 1996), mammals (Moran 1953; Stenseth et al. 1998; Haydon et al. 2001) and birds (Ranta et al. 1995; Koenig 1999; Paradis et al. 1999; Toms et al. 2005), documenting that population dynamics is a regional process in many continuously distributed species.

In the present study I constructed a clustering routine that uses this regional synchronisation to delineate the geographical distributions of populations with synchronised dynamics within, and more desynchronised dynamics across the boundaries of the identified populations. Analysing the Breeding Bird Survey of North American Birds, the clustering routine identified two to five populations per species in 160 species, while no population differentiation was identified for 139 species.

The clustering routine will only aggregate areas that are geographically connected to, and dynamically synchronised with, explicitly identified local synchronisation optima. It requires the presence of at least two local synchronisation optima before it will split a continuously distributed species into more than a single population. It will then delineate the areas that are synchronised with the optima, leaving the population dynamic status and level of synchronisation unresolved for areas that are not synchronised with an explicitly identified local synchronisation optimum.

The population dynamic clustering routine identified fewer population units than the demographic clustering routine of Rushing et al. (2016). For six species of North American passerines, they identified eight to 20 demographic units with an average of 13. For the same eight species, I identified only one to four population dynamic units, with an average of 1.4. One reason for the difference is that Rushing et al. (2016) allowed their method to operate on the scale of single counting routes, while I optimised my routine to operate on the spatial scale that produced the best signal-to-noise ratio for the estimated index trajectories. This scale used grid-squares that were larger than the scale of many of the demographic units identified by Rushing et al. (2016). Also, the inclusion of the mean abundance on counting routes in the clustering routine of Rushing et al. (2016) can be potentially misleading, as variation in mean abundance may simply reflect variation in the relative availability of the relevant habitat across counting routes, identifying clusters that are invariant with respect to the population structure of a species. A third difference is that the routine of Rushing et al. (2016) identifies a statistical number of clusters that differ somewhat from one another. I chose instead to delineate only the population dynamic units that could be explicitly linked to a geographical optimum in population dynamic synchronisation.

These local synchronisation optima may be seen as population dynamics hot-spots that drive the geographic population dynamics of many species. The developed clustering routine includes the surrounding areas with synchronised dynamics in the population components of the optima, until the synchronisation deteriorates at the boundaries between populations that are driven by different optima. These boundaries between different population dynamics can be seen as ecological barriers that surround population dynamic populations, as envisioned conceptually by Andrewartha and Birch (1954) and developed as a statistical data-driven measure of population structure in the present paper.

## Supporting information

SI2: Estimated population trajectories

## 6 Supplementary Information

**Si1:** Trajectory and distribution plots;

**Si2:** Estimated population trajectories;

## Acknowledgements

I thank all that have collected and published the population data I use.

---

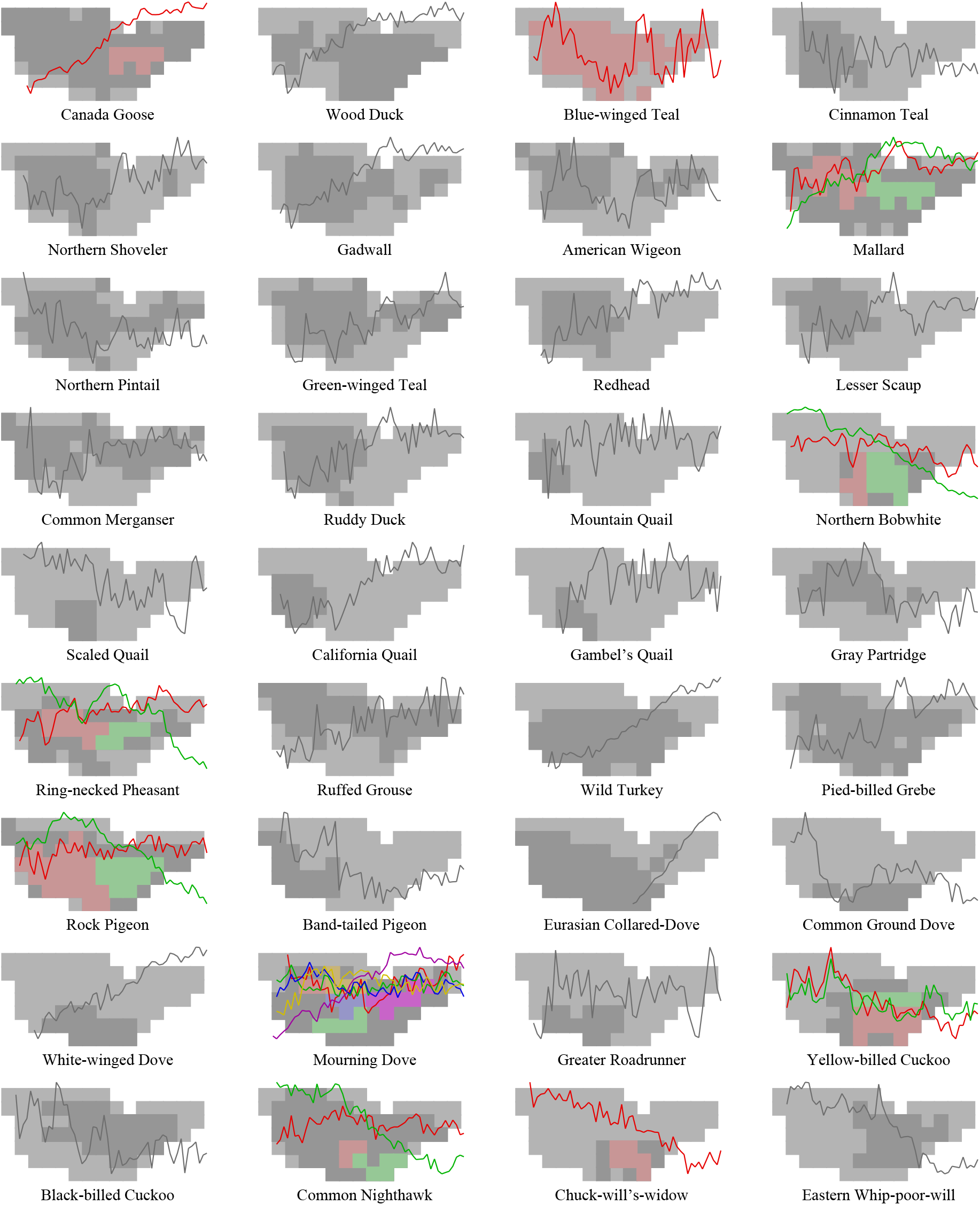

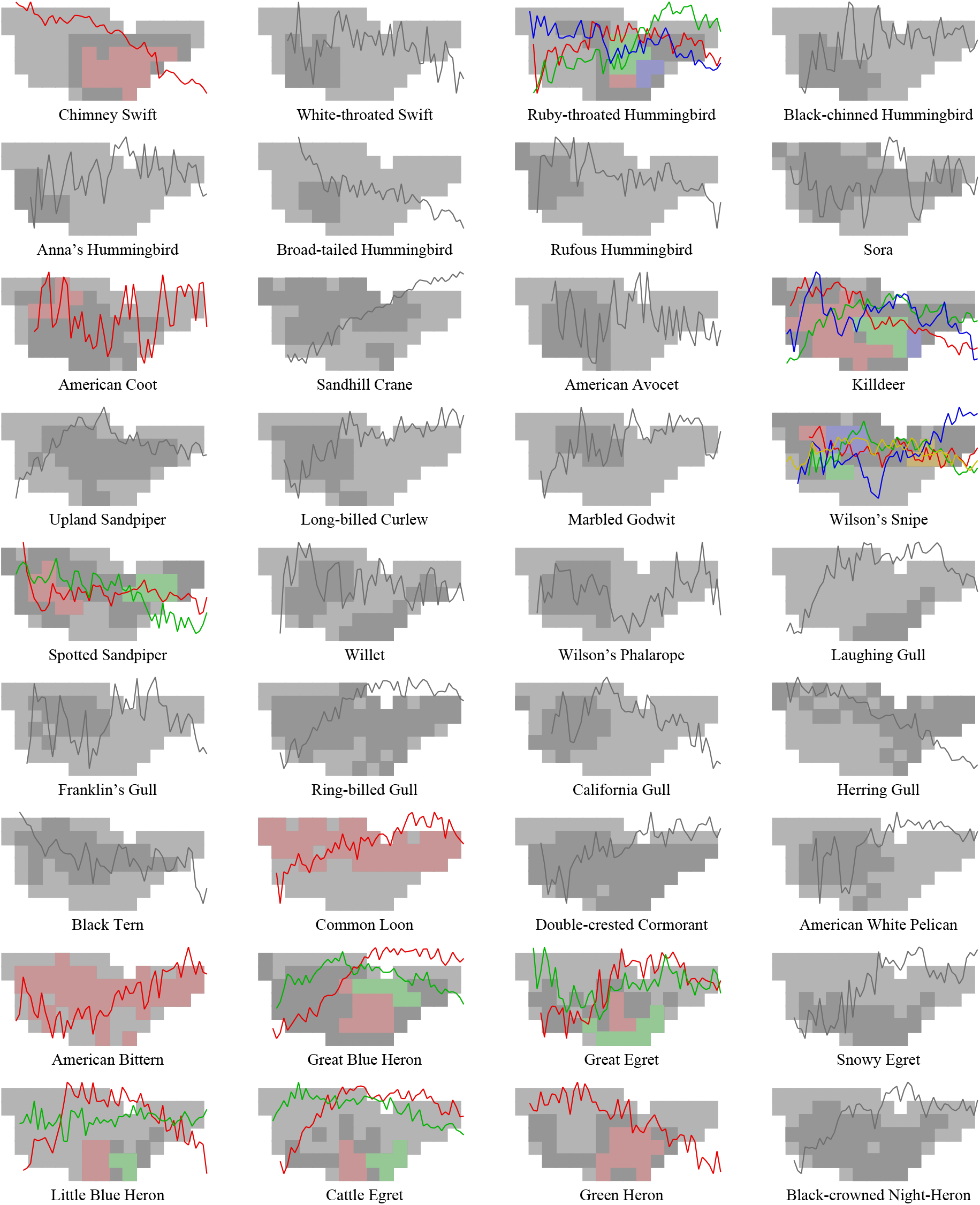

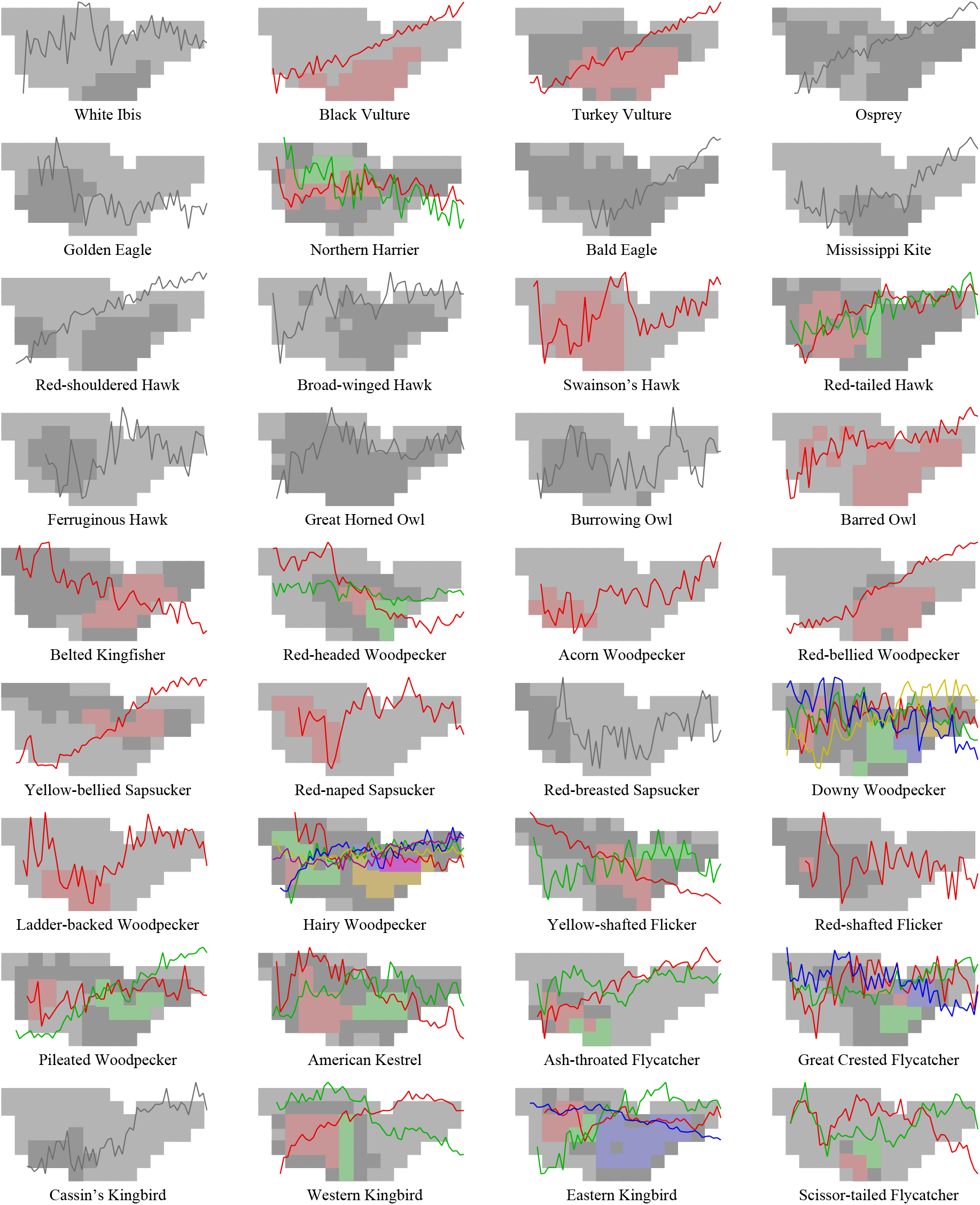

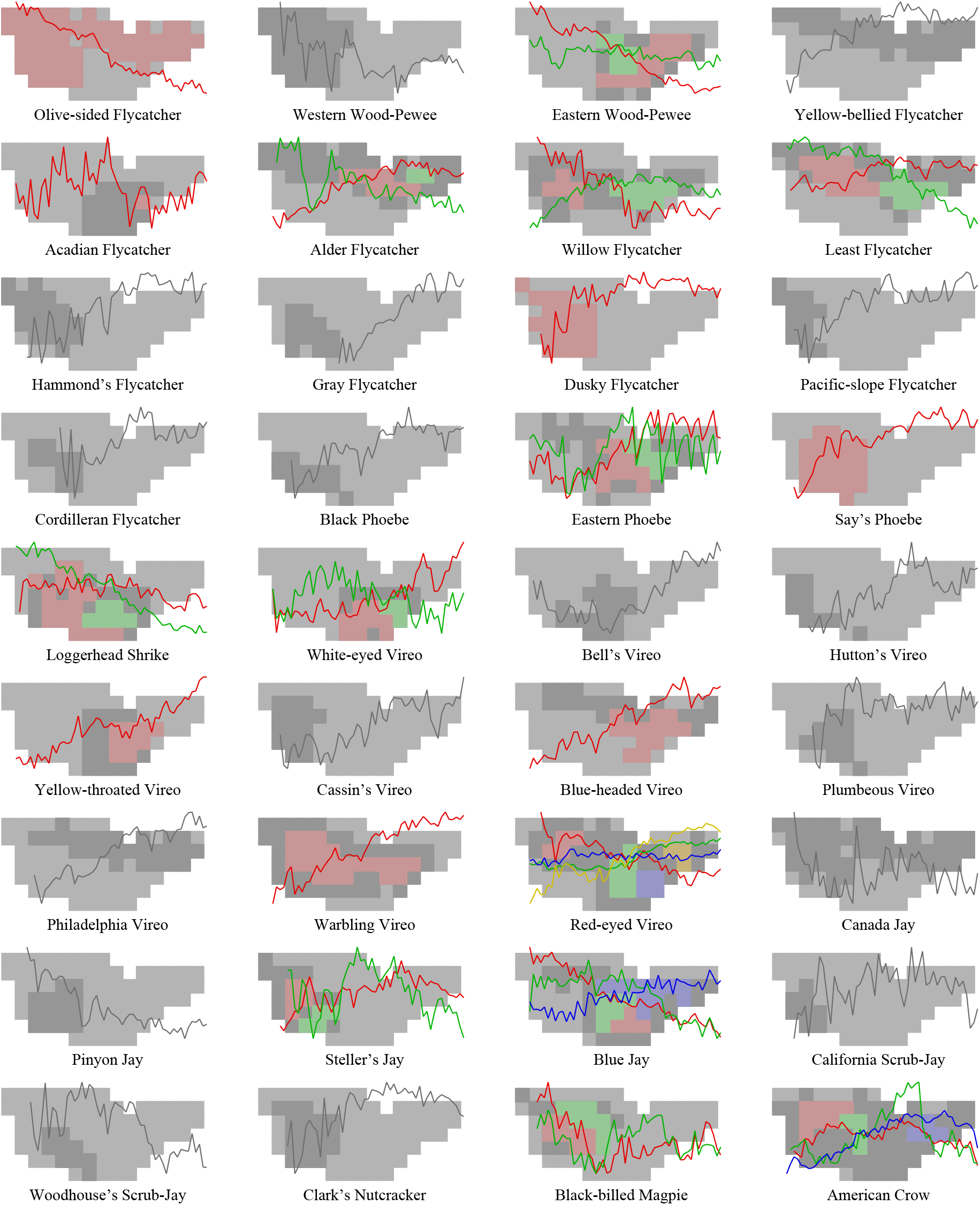

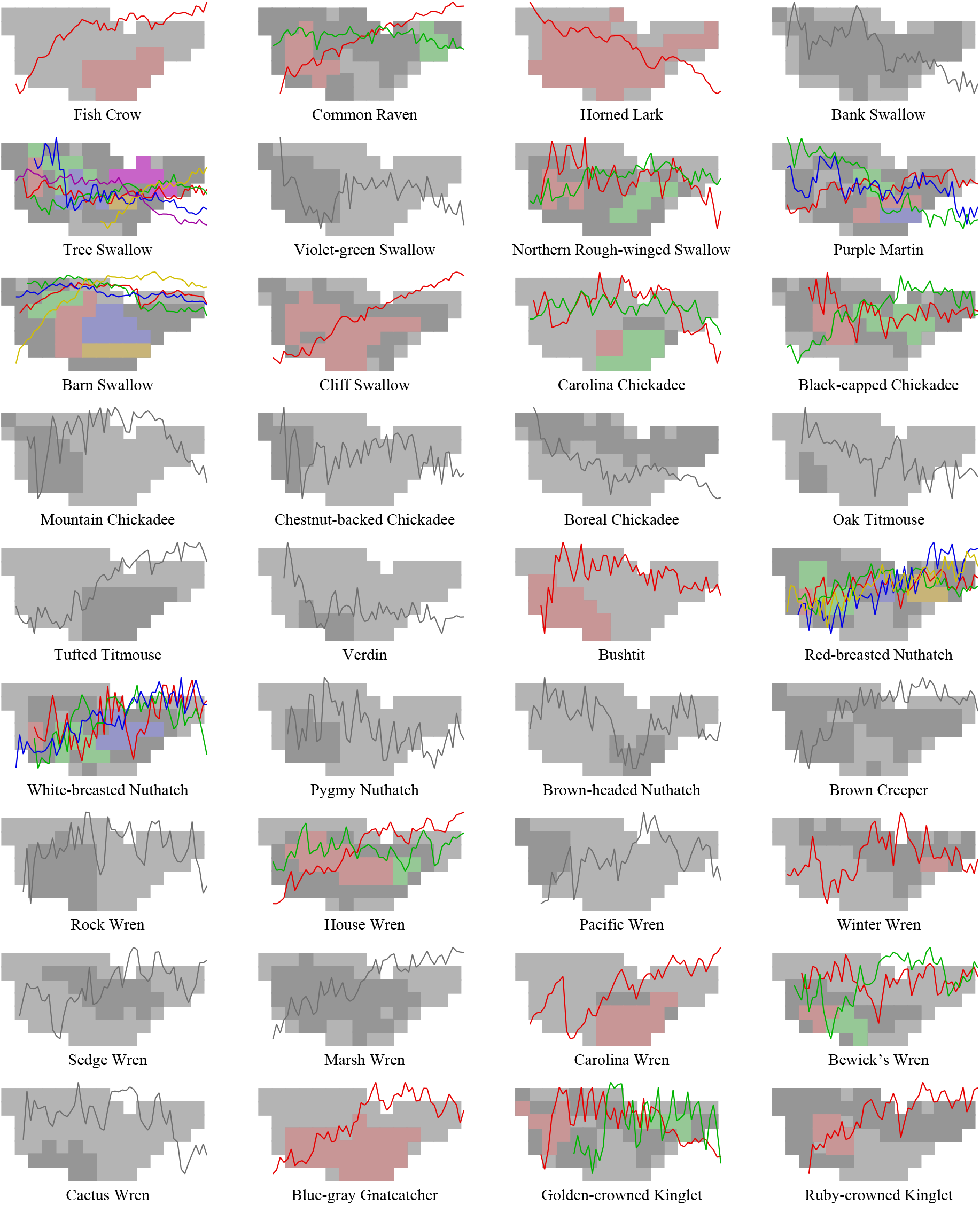

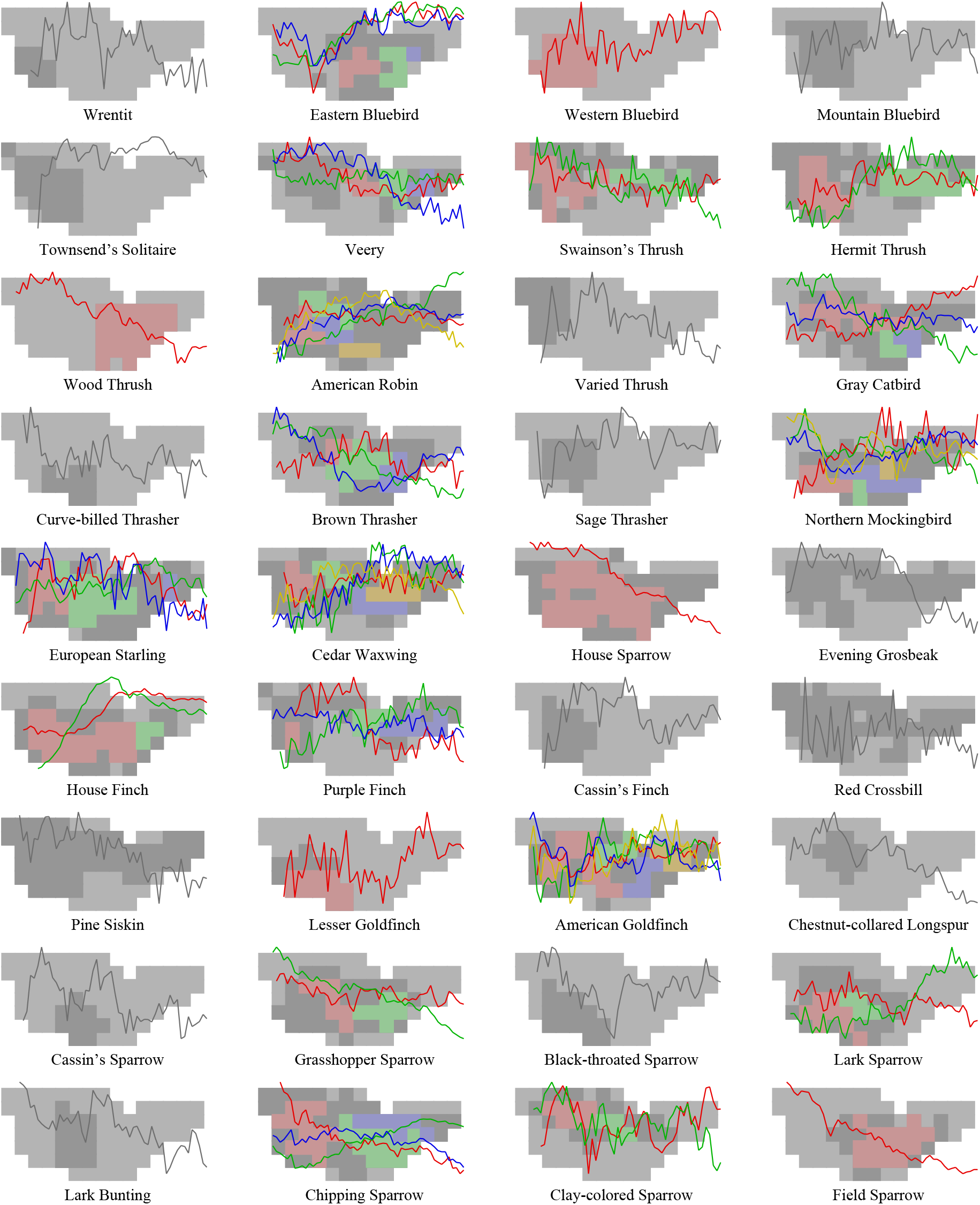

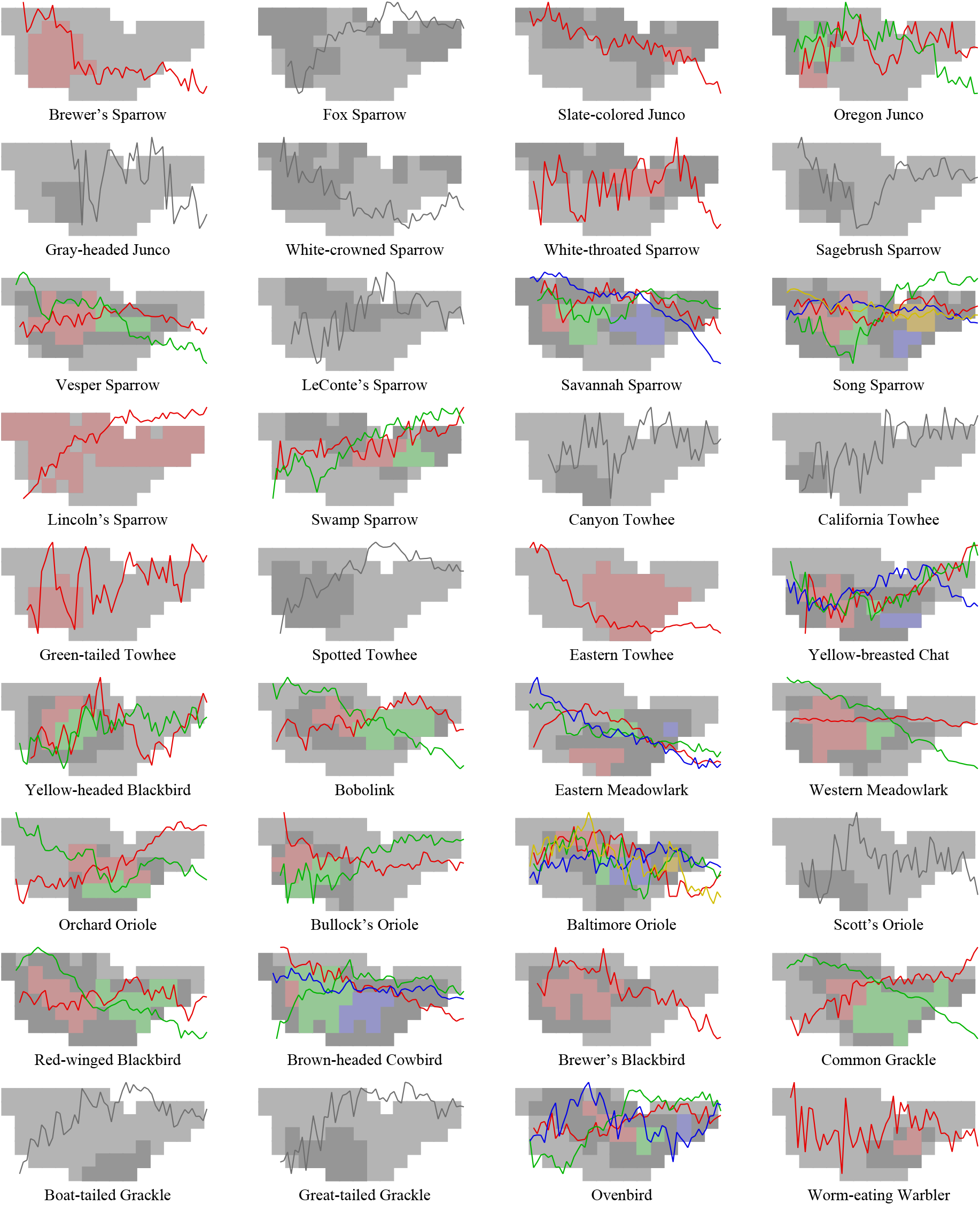

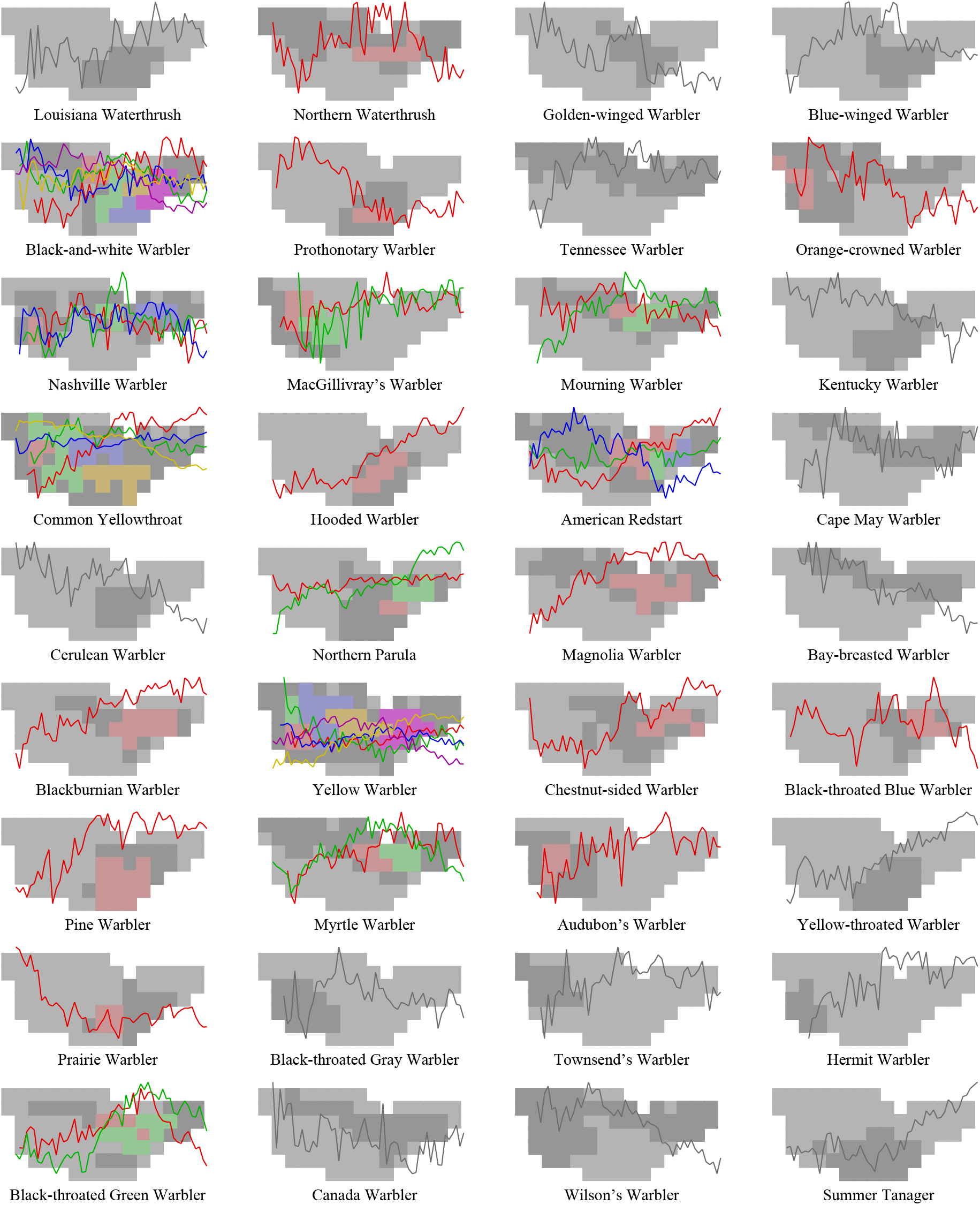

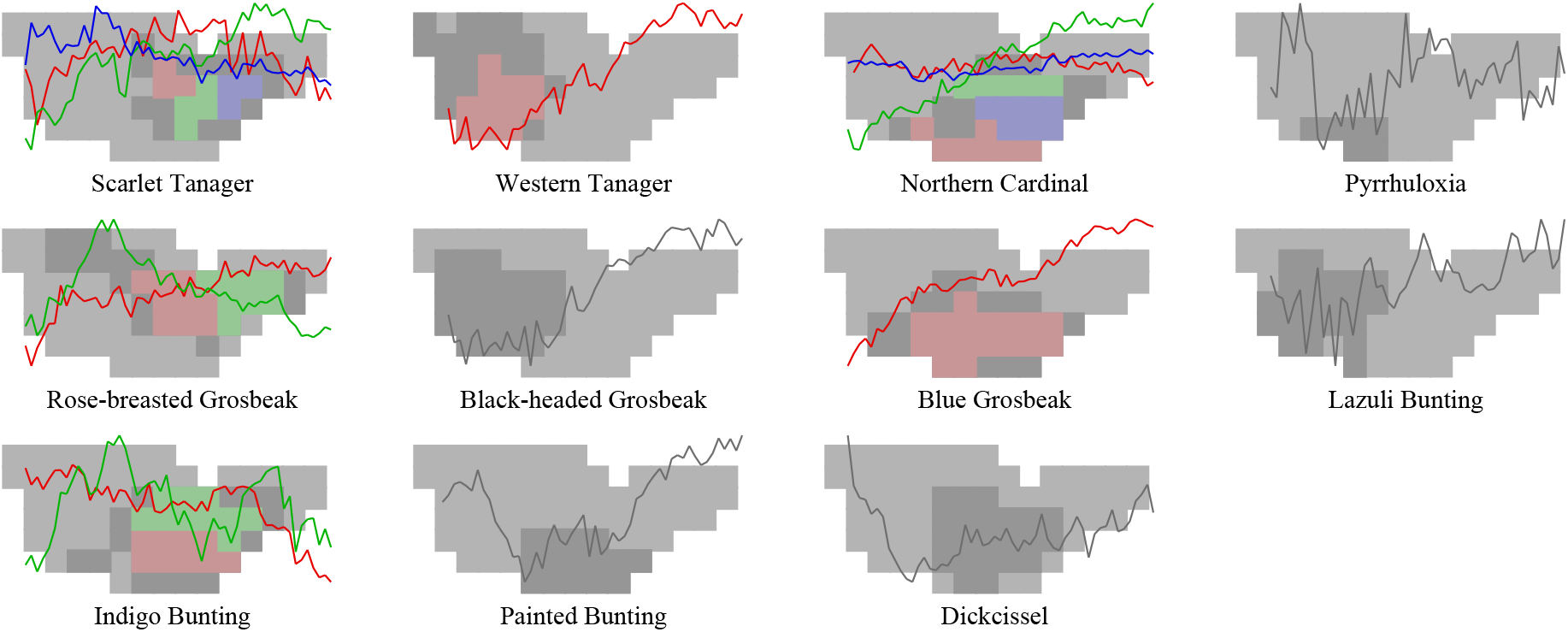

